# Gene Coregulation and Coexpression in the Aryl Hydrocarbon Receptor-mediated Transcriptional Regulatory Network in the Mouse Liver

**DOI:** 10.1101/260018

**Authors:** Navya Josyula, Melvin E. Andersen, Norbert Kaminski, Edward Dere, Timothy R. Zacharewski, Sudin Bhattacharya

## Abstract

Tissue-specific network models of chemical-induced gene perturbation can improve our mechanistic understanding of the intracellular events leading to adverse health effects resulting from chemical exposure. The aryl hydrocarbon receptor (AHR) is a ligand-inducible transcription factor (TF) that activates a battery of genes and produces a variety of species-specific adverse effects in response to the potent and persistent environmental contaminant 2,3,7,8-tetrachlorodibenzo-p-dioxin (TCDD). Here we assemble a global map of the AHR gene regulatory network in TCDD-treated mouse liver from a combination of previously published gene expression and genome-wide TF binding data sets. Using Kohonen self-organizing maps and subspace clustering, we show that genes co-regulated by common upstream TFs in the AHR network exhibit a pattern of co-expression. Specifically, directly-bound, indirectly-bound and non-genomic AHR target genes exhibit distinct patterns of gene expression, with the directly bound targets generally associated with highest median expression. Further, among the directly bound AHR target genes, the expression level increases with the number of AHR binding sites in the proximal promoter regions. Finally, we show that co-regulated genes in the AHR network activate distinct groups of downstream biological processes, with AHR-bound target genes enriched for metabolic processes and enrichment of immune responses among AHR-unbound target genes, likely reflecting infiltration of immune cells into the mouse liver upon TCDD treatment. This work describes an approach to the reconstruction and analysis of transcriptional regulatory cascades underlying cellular stress response using bioinformatic and statistical tools.

## Introduction

“Toxicity pathways” are normal intracellular signaling pathways that when sufficiently perturbed by exogenous chemicals can lead to an adverse outcome at the cellular level, and potentially at the level of tissues and the whole organism (NRC 2007; Whelan and Andersen 2013). Signaling, transcriptional and post-transcriptional regulatory networks underlie toxicity pathways and their dynamic behavior in response to endogenous and exogenous perturbation. It is crucial to understand the organization, structure and dynamics of these networks through mapping and modeling approaches for a quantitative assessment of the risks of chemical exposure to biological systems. Tissue-specific network models of chemical-induced perturbation can improve our understanding of the intracellular events leading to adverse effects and eventual injury from chemical exposure.

The major cellular response pathways are governed both transcriptionally and post-translationally. A core set of master regulatory transcription factors (TFs) are central actors in most molecular pathways leading to altered expression of suites of genes in response to exposure to a variety of chemical compounds (Jennings et al. 2013). These TFs, including the nuclear receptors, p53, nuclear factor erythroid 2-related factor (Nrf2), nuclear factor-κB (NF-κB), the STAT (signal transducers and activators of transcription) family and the aryl hydrocarbon receptor (AHR), typically coordinate a broad range of physiological processes like metabolism, oxidative stress response, differentiation, tumor suppression, reproduction, development and homeostasis (Tyagi et al. 2011; Ma 2013; Evans and Mangelsdorf 2014; Audet-Walsh and Giguere 2015; Wright et al. 2017). They thus act as sentinels of normal biological activity, but their inappropriate activation or inhibition can lead to adverse outcomes at the cellular or tissue level (Andersen et al. 2013).

Here we describe a network model of the AHR pathway in the mouse liver, assembled from previously published genomic data sets and newly analyzed using various computational methods. The AHR is a ligand-activated TF that belongs to the basic helix-loop-helix (bHLH)-PER-ARNT-SIM (PAS) family of proteins, which serve as sensors of developmental and environmental signals (Gu et al. 2000). The prototypical AHR ligand is TCDD (Poland et al. 1976), a persistent environmental toxicant that produces a variety of adverse effects in laboratory animals, including immune suppression, reproductive and endocrine effects, neurochemical alterations, developmental toxicity, chloracne and tumor promotion (Birnbaum 1994; Pohjanvirta and Tuomisto 1994). These effects are mediated by the transcriptional activity of the AHR, as shown by their absence or amelioration in AHR-null mice and mice with low-affinity AHR alleles (Okey et al. 1989; Gonzalez and Fernandez-Salguero 1998; Peters et al. 1999), as well as in mice with mutations in the DNA-binding domain or nuclear localization sequence of the AHR (Bunger et al. 2003; Bunger et al. 2008). Ligand binding causes the inactive AHR in the cytosol to undergo a conformational change resulting in dissociation from its chaperone protein complex and translocation to the nucleus, where it forms a heterodimer with the related nuclear protein aryl hydrocarbon nuclear translocator (ARNT) (Hoffman et al. 1991; Whitelaw et al. 1993). The AHR-ARNT complex then binds to specific DNA sequences on target genes called dioxin response elements (DRE) containing the core sequence 5’-GCGTG-3’ (Denison et al. 1988), leading to the regulation of a diverse battery of genes (Poland and Knutson 1982; Hankinson 1995). While the 5’-GCGTG-3’ nucleotide core is substitution-intolerant, the flanking 5’ and 3’ nucleotides adjacent to the core sequence also contribute to a functional AHR binding site (Denison et al. 1988; Shen and Whitlock Jr 1992; Lusska et al. 1993; Gillesby et al. 1997). DRE-independent mechanisms of AHR binding have also been reported (Dere et al. 2011b; Huang and Elferink 2012).

While the density of AHR-bound regions in the genome of hepatic tissue from TCDD-treated mice is greatest in proximal promoter regions close to the transcription start site (TSS) of annotated genes, AHR also binds to sites distal from a TSS, e.g. in intergenic regions and 3’ UTRs (Dere et al. 2011b). Moreover, only a third of the differentially expressed genes identified by microarray analysis showed AHR binding at a DRE in their proximal promoter regions, suggesting additional mechanisms of gene regulation by AHR beyond the canonical model described above (Dere et al. 2011b). These mechanisms may include target gene regulation from distal AHR-bound regions through DNA looping, or indirect regulation by AHR through tethering with a secondary TF (Farnham 2009). Such an indirect mechanism has been demonstrated in the regulation of the rat CYP1A2 gene by AHR (Sogawa et al. 2004).

Here we have mapped the TCDD-induced AHR regulatory network from a combination of gene expression and ChIP-on-chip data from the mouse liver (Dere et al. 2011b), which provides us a system-wide view of AHR-mediated gene regulation under short-term TCDD exposure (168 hr). Specifically, statistical and visualization tools were used to establish a relationship between gene co-regulation by multiple TFs and gene co-expression, and link groups of co-regulated genes to distinct downstream functional outcomes. Our focus here is on the early stages of hepatic response to TCDD exposure - longer-term exposure may lead to a different suite of adaptive responses at the cellular and tissue level.

## Methods

### Microarray data

Our network analysis was based on results from a previous study of gene expression profiling using whole genome oligonucleotide arrays (Agilent Technologies, Santa Clara, CA) of hepatic tissues from female C57BL/6 mice orally gavaged with 30 μg/kg of TCDD (Boverhof et al. 2005; Dere et al. 2011b). The gene expression analysis was performed in hepatic tissue from mice exposed to TCDD for 2, 4, 8, 12, 18, 24, 72 and 168 hrs. Differentially responsive genes were identified using previously described cutoffs for fold change and statistical significance (fold change | ≥ 1.5 and posterior probabilities P1 (t) ≥ 0.999) (Eckel et al. 2004; Dere et al. 2011b).

### ChlP-on-chip data

Genome-wide AHR location data were taken from the previously described ChIP-on-chip experiments (Dere et al. 2011b), where ChIP assays were performed with hepatic tissue from female C57BL/6 mice exposed to TCDD for 2 and 24 hrs. Genes were associated with AHR-enriched regions if the position of maximum fold enrichment was within 10 kb upstream of a transcriptional start site (TSS) through to the end of the 3’ UTR. For the present analysis, the ChIP data for 2 and 24 hrs. were combined to obtain a unique list of ChIP enriched regions associated with annotated genes (Supplementary Methods; Supplementary Code 1).

### DRE analysis in ChIP enriched regions

The ChIP enriched regions for the differentially expressed (DE) genes were computationally searched for the presence of 5’-GCGTG-3’ DRE core sequences to infer the nature of AHR binding to the target genes. The putative DRE search algorithm, written in *R* (R Core Team 2016) (Supplementary Methods; Supplementary Code 2), was based on a previously described approach (Sun et al. 2004). Briefly, the genomic sequences of the enriched regions were obtained from UCSC Genome Browser (http://genome.ucsc.edu) and scanned for exact matches to the DRE core sequences on both positive and negative strands. For each matched region, the 5-bp core sequence was extended 7 bp upstream and downstream of the core. The matrix similarity (MS) scores (Quandt et al. 1995) for the 19-bp DRE sequences were calculated and compared to an MS score threshold of 0.8473 based on the lowest MS score of 13 bona fide AHR-binding sequences (Dere et al. 2011a) (i.e. sites from the literature confirmed to bind AHR). The DRE sequences with high MS scores (MS score ≥ 0.8473) were defined as putative DREs capable of binding AHR. The DE genes that were AHR-enriched and had a putative DRE in the enriched region were described as “directly bound” by AHR, while AHR-enriched genes without a putative DRE were described as “indirectly bound”. The remaining DE genes that were not AHR-enriched were regarded as “unbound” / “non-genomic” targets.

### Construction and visualization of the AHR transcriptional regulatory network

The DE genes from the Agilent oligonucleotide array data were searched against online databases to obtain a list of TFs that regulate these genes. The ChIP-X Enrichment Analysis (ChEA2) database (Kou et al. 2013) was used to obtain the list of regulatory TFs. To obtain the mouse-liver specific list of transcription factors, the mouse-specific TFs from ChEA2 were screened for expression in the liver using the TRANSFAC^®^ database (Matys et al. 2003). The ensemble of DE genes including the directly and indirectly AHR-bound genes, together with their inferred transcriptional regulators, form a comprehensive network for TF-gene interactions under AHR-mediated TCDD induction. The landscape of this regulatory network was rendered using the open source network visualization tool Cytoscape (Shannon et al. 2003). The gene expression values at each time point of TCDD exposure were superposed on this network to visualize the temporal changes associated with each gene. A | fold ratio | threshold of 1.20 was used to identify the key target genes that are themselves TFs regulating other genes in the dataset (a looser fold change threshold was used for TFs than other genes as TFs tend to be more tightly regulated). To generate and annotate the network in Cytoscape, three input files describing the network topology and gene expression values were used: an AHR-gene interaction file and a TF-gene interaction file (“network files”), and a gene expression file (“attributes file”). Log2 scaling of the fold ratios was used for visualizing gene expression. The network files were merged together to form the complete layout.

### Gene expression analysis based on transcriptional groupings

A binary TF-gene interaction matrix with 43 TFs in addition to AHR was created indicating which TFs interact with which target genes. If a gene is regulated by a particular TF, then the corresponding interaction is represented as ‘1’; otherwise it is represented as ‘0’. We used this TF-gene interaction matrix to classify target genes into co-regulated groups in a transcriptional cascade, in order to examine any possible relation between co-regulation and co-expression. To generate this grouping, AHR and other key TFs that were also target genes were considered in all possible combinations to identify the expression trends for target genes in each group. The total number of genes in each co-regulated group was counted by referring to the TF-gene interaction matrix, and all groups with at least 5 genes were considered for examination of the expression patterns. A graphical analysis was performed in *R* to identify the expression patterns of target genes for each combination of regulatory TFs (Supplementary Methods; Supplementary Code 3).

### Kohonen self-organizing maps to visualize gene co-expression

To further examine the relationship between the transcriptional groups and target gene expression patterns, a self-organizing map (SOM) for the AHR network was generated using the Kohonen SOM package in *R* (Wehrens and Buydens 2007). The same TF-gene interaction matrix described above was used as input for this analysis. The SOM algorithm follows a clustering technique to group the target genes according to their TF binding patterns. Target genes with similar TF binding patterns are grouped into the same cluster or adjacent clusters, referred to as ‘units’ (Supplementary Methods; Supplementary Code 4).

### Subspace clustering

The ORCLUS subspace clustering algorithm (Aggarwal and Yu 2000) and corresponding *R* package (Szepannek 2013) were used to cluster the differentially expressed genes into 16 non-overlapping groups. The number of clusters *k* = 16 and the dimensionality of each cluster *l* = 4 were chosen so as to minimize the cluster sparsity coefficient (Aggarwal and Yu 2000) (**Supplementary Code 5**).

### Functional categorization of genes in each cluster

Gene ontology (GO) functional analysis was performed for the DE genes present in each ORCLUS cluster. Enriched GO “process” categories were identified for genes in each cluster using the *GOrilla* tool (Eden et al. 2009) with a p-value threshold of 10^−3^ and the list of all DE genes as background. *REViGO* (Supek et al. 2011) was used to arrange the enriched processes into a “treemap”, which was then rendered as an image using the downloadable R script generated by the program (**Supplementary Code 6, Supplementary Code 7**).

## Results

### Differential gene expression

The raw array dataset (Dere et al. 2011b) consisted of 41,267 records with annotated genes, fold ratio and significance (P1 (t) values) at 2, 4, 8, 12, 18, 24, 72 and 168 hrs. post TCDD exposure. For genes with multiple occurrences in the dataset, the fold ratios and P1(t) values were averaged, resulting in a total of 21,307 unique gene records. After applying the statistical cutoff values for fold change and P1(t) at each expression time point, the resulting number of unique differentially expressed (DE) genes was 1,407. All 1,407 DE genes were used to generate the AHR regulatory network map.

### Analysis of AHR-enriched genomic regions associated with DE genes

The ChlP-on-chip datasets for 2 and 24 hrs. time points consisted of 14,446 and 974 AHR-enriched regions respectively, with associated genes (Dere et al. 2011b). The two datasets were combined to yield a unique list of genes associated with at least one enriched region. This list of enriched genes was compared against the list of 1,407 DE genes, yielding 632 genes associated with one or more AHR-enriched regions. The AHR-enriched regions around these 632 genes were searched for putative DREs, producing three kinds of regions depending on presence and location of DREs:

a. Regions with one or more 5-bp DRE cores centrally located such that a 7-bp upstream and downstream extension was possible for MS score calculations.
b. Regions with DRE cores present only at the edge of the region so that the 7-bp extension in both directions was not possible.
c. Regions with no DRE core.

A total of 144 genes were associated with AHR-enriched regions where MS score calculations were possible, and that had putative DREs, i.e., 19-bp DRE sequences with an MS score ≥ 0.8473 (see Methods). These genes were considered to be “directly bound” by AHR. For the AHR-enriched regions with (i) non-putative DRE core (i.e. MS score < 0.8473), (ii) DRE core located at edges, or (iii) DRE core not present in the enriched region, the associated genes were considered to be “indirectly bound” by AHR. In total, among the 1,407 differentially expressed genes, 632 were bound by AHR with 144 genes directly bound, 488 indirectly bound, and the remaining 775 genes unbound by AHR.

### Other transcriptional regulators of the DE genes

The ChEA2 database (Kou et al. 2013) provides a comprehensive record of transcription factor/target gene interactions from genome-wide ChIP studies for both mouse and human. The list of 1,407 DE genes from our analysis was uploaded to the ChEA2 server, which identified 104 unique mouse-specific transcriptional regulators for these genes. These 104 TFs were searched against the TRANSFAC^®^ database to filter for expression in liver tissue, which after accounting for discrepancies in naming between ChEA2 and TRANSFAC^®^ resulted in a list of 43 unique mouse liver-expressed TFs. Out of our 1,407 DE genes, 1,198 had interactions with at least one of these 43 transcription factors. Among the 43 TFs, seven were themselves target genes of other identified TFs differentially expressed at |fold ratio| > 1.2 in the microarray dataset. These seven TFs were NRF2, FLI1, KLF4, SOX17, CCND1, PPARG and GATA1, and in addition to AHR, form the “hubs” of the inferred mouse liver AHR network.

### The AHR regulatory network

All interactions of the DE genes with AHR and the other 43 identified TFs together form the mouse liver AHR regulatory network (**Figure 1**), which consists of 44 “source” nodes interacting with 1,241 “target” nodes.

**Figure 1:**
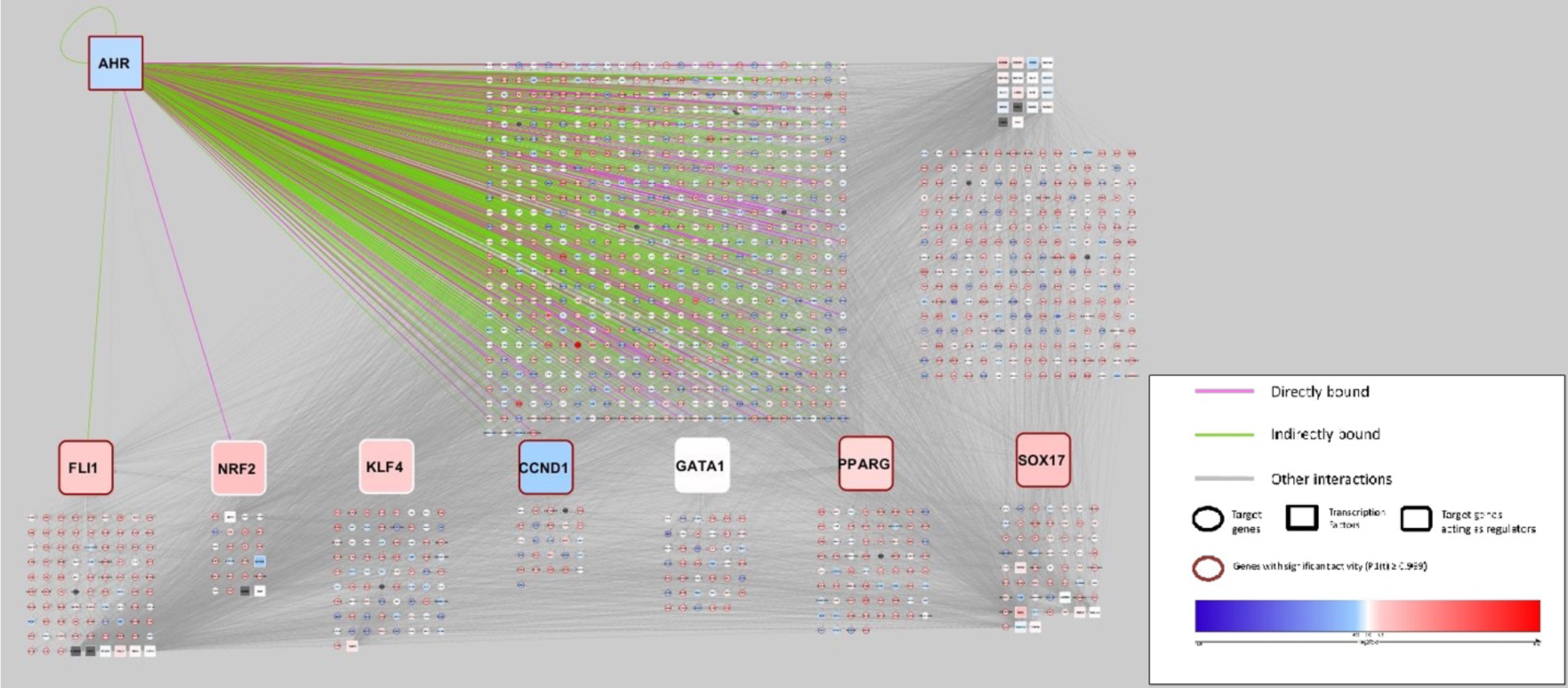
A comprehensive AHR transcriptional regulatory landscape in the mouse liver, viewed at time t = 168h and TCDD dose = 30 μg/kg. The AHR network in the mouse liver reveals a hierarchical structure. Transcription factor (TF) nodes are shown as rectangles, target genes as circles. Edges represent TF-target gene binding. AHR is at the top left of the figure, with AHR interactions shown with pink and green edges indicating direct and indirect binding respectively. The seven key non-AHR TFs that are also target genes of other TFs are shown in the bottom half of the figure along with their target genes. Other TFs in the network and their target genes are shown in the top right of the figure. Each node is colored according to their log2 fold change ratio at 168hrs (nodes in grey were not differentially expressed at 168hrs). Non-AHR TF-gene interactions are shown as grey edges.

AHR and the other seven hub TFs act as both source and target nodes (AHR regulates itself). Two of these hub TFs are regulated by AHR: Nrf2 is a direct target and Fl i 1 an indirect target (**Figure 1**). The expression levels for up- and down-regulated genes were superposed on this network layout for each of the eight time points in the gene array study (**Supplementary Figure 1a-h**), illustrating that the gene expression levels were not monotonic in time.

To examine the transcriptional regulatory hierarchy in the network, AHR and four of the seven hub TFs (Fli1, Nrf2, Klf4 and Sox17), which were all expressed at |fold ratio | > 1.5, were grouped in all possible combinations (**Supplementary Table 1**) to assess expression of their target genes. Genes differentially expressed at least at one time point at | fold ratio | > 1.5 were chosen for this analysis, yielding 1,191 target genes regulated individually or in combination by the above five TFs (**Supplementary Table 1**). We then examined the expression pattern of groups of co-regulated genes with a count of 5 or more (**Table 1**). The time courses of genes that were up-regulated at the 168 hrs. time point (**Figure 2**) suggest that genes with the same upstream regulators have similar expression patterns.

**Figure 2:**
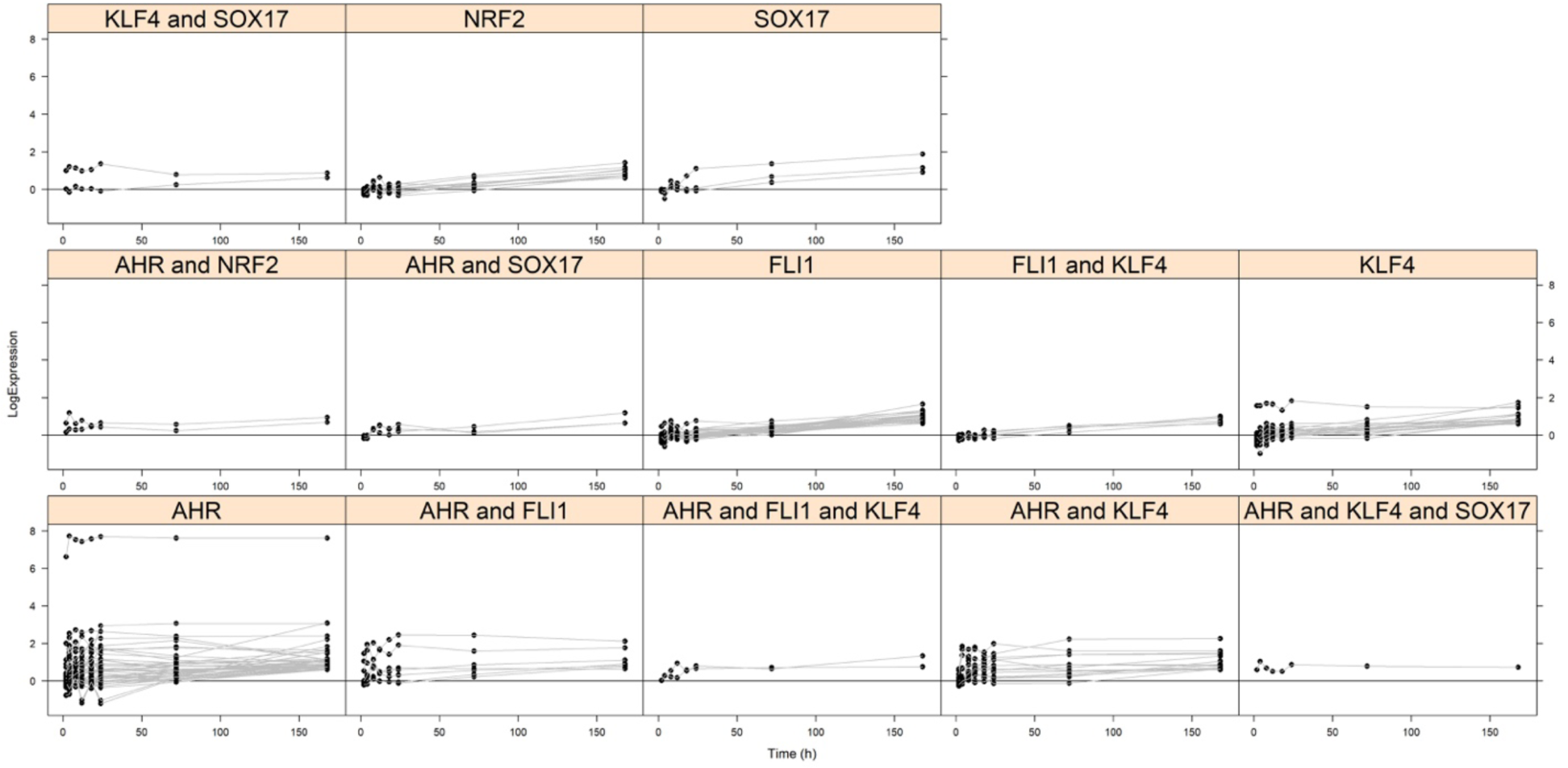
Time courses of genes grouped by transcriptional regulators (only genes up-regulated at 168hrs. shown). Genes grouped by transcriptional regulators show similar expression patterns. The vertical axis denotes log2 fold change.

**Table 1:** Count of genes in each transcriptional grouping. All possible groupings among the “hub” TFs in the network, AHR, Flil, Nrf2 (NFe2L2), Klf4 and Soxl7 were considered. Groups with gene counts > 5 are shown in the table.

|  | Combinations | TF.Groups | Gene.Count |
| --- | --- | --- | --- |
| 1 | AHR | G1 | 210 |
| 2 | FLI1 | G2 | 36 |
| 3 | NFE2L2 | G3 | 13 |
| 4 | KLF4 | G4 | 55 |
| 5 | SOX17 | G5 | 10 |
| 6 | AHR\|FLI1 | G6 | 15 |
| 7 | AHR\|NFE2L2 | G7 | 12 |
| 8 | AHR\|KLF4 | G8 | 54 |
| 9 | AHR\|SOX17 | G9 | 7 |
| 10 | FLI1\|KLF4 | G11 | 9 |
| 11 | KLF4\|SOX17 | G15 | 6 |
| 12 | AHR\|FLI1\|KLF4 | G17 | 7 |
| 13 | AHR\|KLF4\|SOX17 | G21 | 5 |

### Co-regulation and co-expression in the AHR network

To take a closer look at whether co-regulation in the AHR network is associated with co-expression, we clustered the 1191 target genes into self-organizing maps (SOMs) based on the factors that regulate them (**Figure 3A**). Each circular unit in the SOM represents a grouping of genes (individual dots within a unit). The SOM algorithm groups genes into units such that genes in a single unit or adjacent units have a similar combination of TFs regulating them (no gene expression values were used for clustering), while genes in distant units have more dissimilar regulators. The median expression level (log2 fold change) of the genes in each unit was then superposed on the SOM as a continuous color scale with blue indicating suppression and red activation (panels in **Figure 3B**). A distinct pattern emerges over the time course, with the units with high median expression at 168 hrs. localized at the lower right corner of the SOM (**Figure 3B**). The median time courses of the genes in adjacent units are also quite similar (**Supplementary Figure 2**). This analysis shows a strong association between gene co-regulation and co-expression in the AHR network.

**Figure 3:**
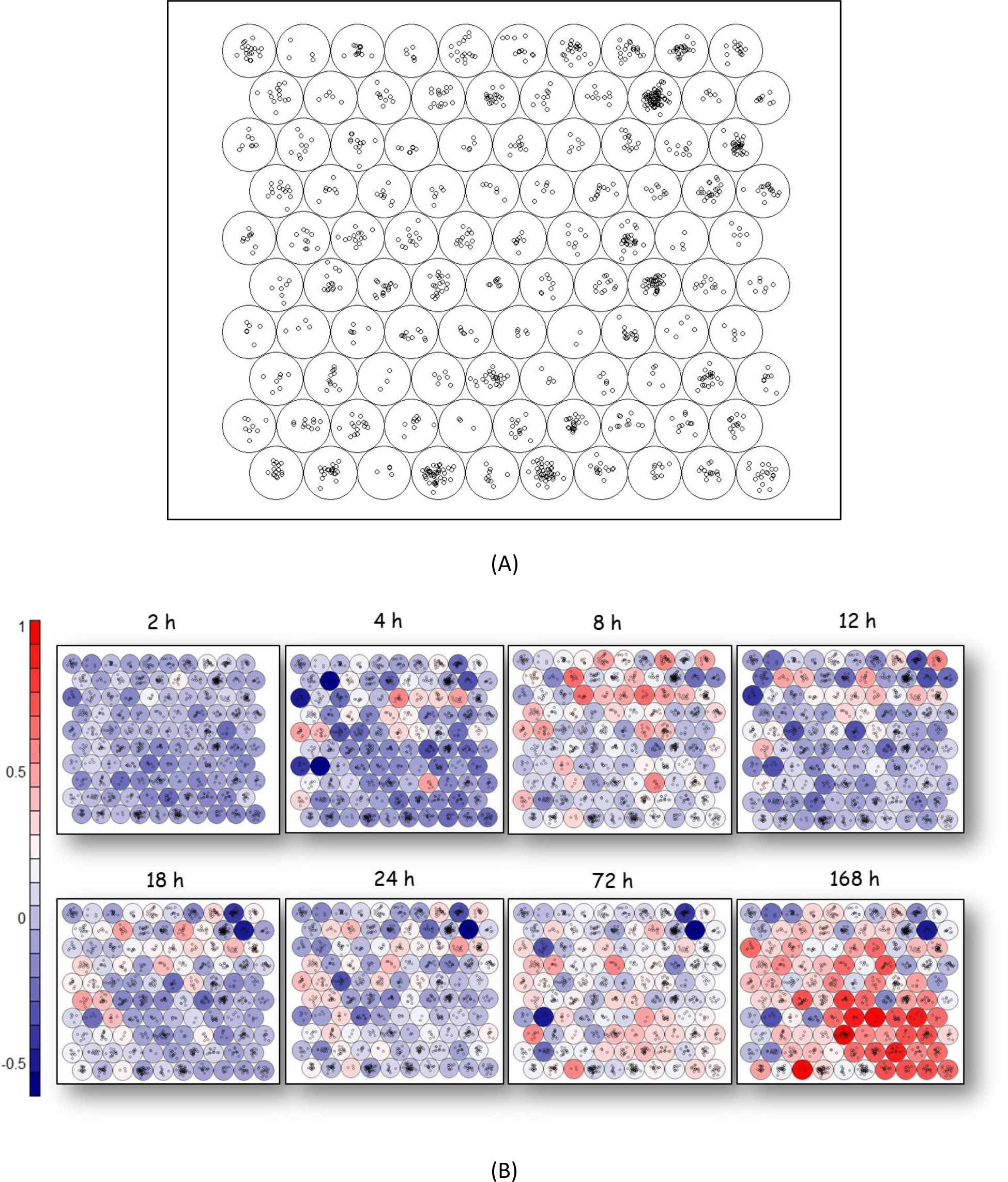
Median expression of genes organized into Kohonen Self-Organizing Map (SOM). (A) The mapping of genes clustered in each unit according to TF binding patterns. (B) The temporal gene expression patterns of the SOM units, confirming the coregulation and coexpression patterns of the genes. The continuous color scale shows the median log_2_ fold change expression values for the genes in each unit, with blue indicating suppression and red activation.

### Localized clustering of co-regulated genes

We further attempted to cluster the 1,191 target genes considered in the SOM analysis above based on regulation by the 44 TFs. Fundamentally, the clustering problem may be stated as: “Given a set of data points, partition them into a set of groups which are as similar as possible” (Aggarwal 2014). If we consider the binary TF-gene connectivity matrix, with genes in rows (observations), TFs in columns (features) and each matrix element equaling 1 or 0 depending on whether a TF binds a gene, we have a high-dimensional clustering problem with feature localization, i.e. different groups of genes are regulated by different subsets of TFs. Global clustering methods like k-means, or dimensionality reduction approaches like principal components analysis do not perform well in this situation, which motivated the development of high-dimensional subspace clustering methods (Aggarwal 2014). These methods include “projected clustering” or “subspace clustering” approaches like PROCLUS (Aggarwal et al. 1999), CLIQUE (Agrawal et al. 2005) and ORCLUS (Aggarwal and Yu 2000), where feature selection or transformation is performed specific to different localities of the data (Aggarwal 2014). ORCLUS in particular is suited for data sets like ours where relevant subspaces may be arbitrarily oriented due to inter-feature correlations (Aggarwal and Yu 2000), i.e. many TFs are correlated in term of which genes they regulate.

We used the ORCLUS algorithm to group the DE genes by TF connectivity into 16 clusters (**Figure 4A**), illustrating both the sparsity of the TF-gene connectivity matrix and the fact that different clusters of genes are regulated by different subsets of TFs. In particular, there is a marked contrast between Cluster 2, where none of 157 genes is bound by AHR, and Cluster 6, where all 123 genes are. Cluster 2 genes can thus be said to comprise a “non-genomic pathway” and Cluster 6 genes a “genomic pathway” with respect to regulation by AHR. The most frequent regulators in Cluster 2 are PPARG, regulating 46 genes, STAT3 (33 genes), CEBPB, NANOG (30 genes each), CREB1, GATA2 and SUZ12 (27 genes each). In contrast, the most frequent regulators in Cluster 6 are AHR (all 123 genes), followed by SUZ12 (55 genes), PPARG (42 genes), CREB1 (29 genes), MYC (25 genes), NANOG (24 genes), E2F1 and TAL1 (22 genes each).

**Figure 4:**
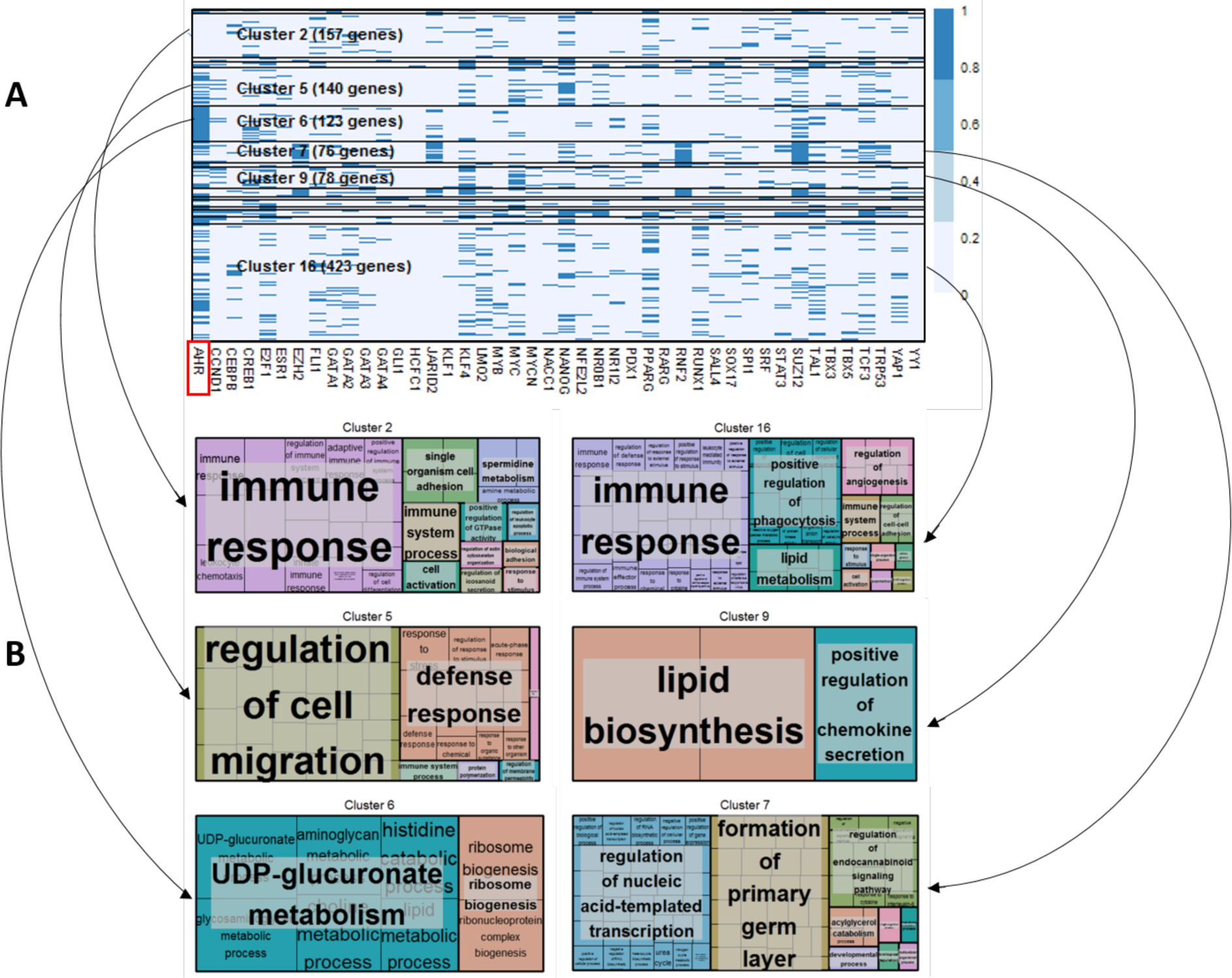
Distinct co-regulated clusters of genes in the AHR network activate distinct downstream pathways. (A) Clustered TF-gene connectivity matrix with 1191 genes in rows and 44 TFs in columns. Elements of the matrix have the value 1 (dark blue) if the corresponding TF and gene are bound; and 0 (pale blue) otherwise. Clusters of more than 50 genes are labeled. (B) Gene Ontology (GO) process categories associated with clusters of more than 50 genes. Sizes of individual boxes representing GO processes are proportional to negative logiQ p-values of enrichment.

### AHR binding and gene expression

We plotted the time courses of log2 fold change values for all genes in Clusters 2 and 6 (**Figure. 5A and 5B**). Genes in the “non-genomic” Cluster 2 are downregulated or moderately upregulated at earlier times (**Figure 5A**), with about two-thirds of the genes showing upregulation at the later time points. On the contrary, a majority of genes in the “genomic” Cluster 6 (**Figure 5B**) are moderately to strongly upregulated at all time points, with a smaller subset showing consistent downregulation. This led us to suspect that there may be a link between binding of a gene by AHR and its expression level, and we separately plotted the time courses of genes that are (a) directly bound, (b) indirectly bound, and (c) unbound by AHR (**Figure. 6A-C**). Nearly all genes in the directly bound group (**Figure 6A**) are upregulated moderately or strongly at multiple time points, whereas in the indirectly bound group (**Figure 6B**), about half of the genes are consistently downregulated. Finally, the unbound group shows an unusual pattern of gene expression (**Figure 6C**), where most genes are downregulated at the first two time points, but then about two-thirds of the genes show progressive upregulation up to the 168 h time point. This observation suggests a cascade structure in the AHR network, where genes not proximally bound by AHR are bound by other TFs activated at intermediate to later points in the time course, or are targets of long range interaction with distally-bound AHR, leading to their upregulation at later times.

**Figure 5:**
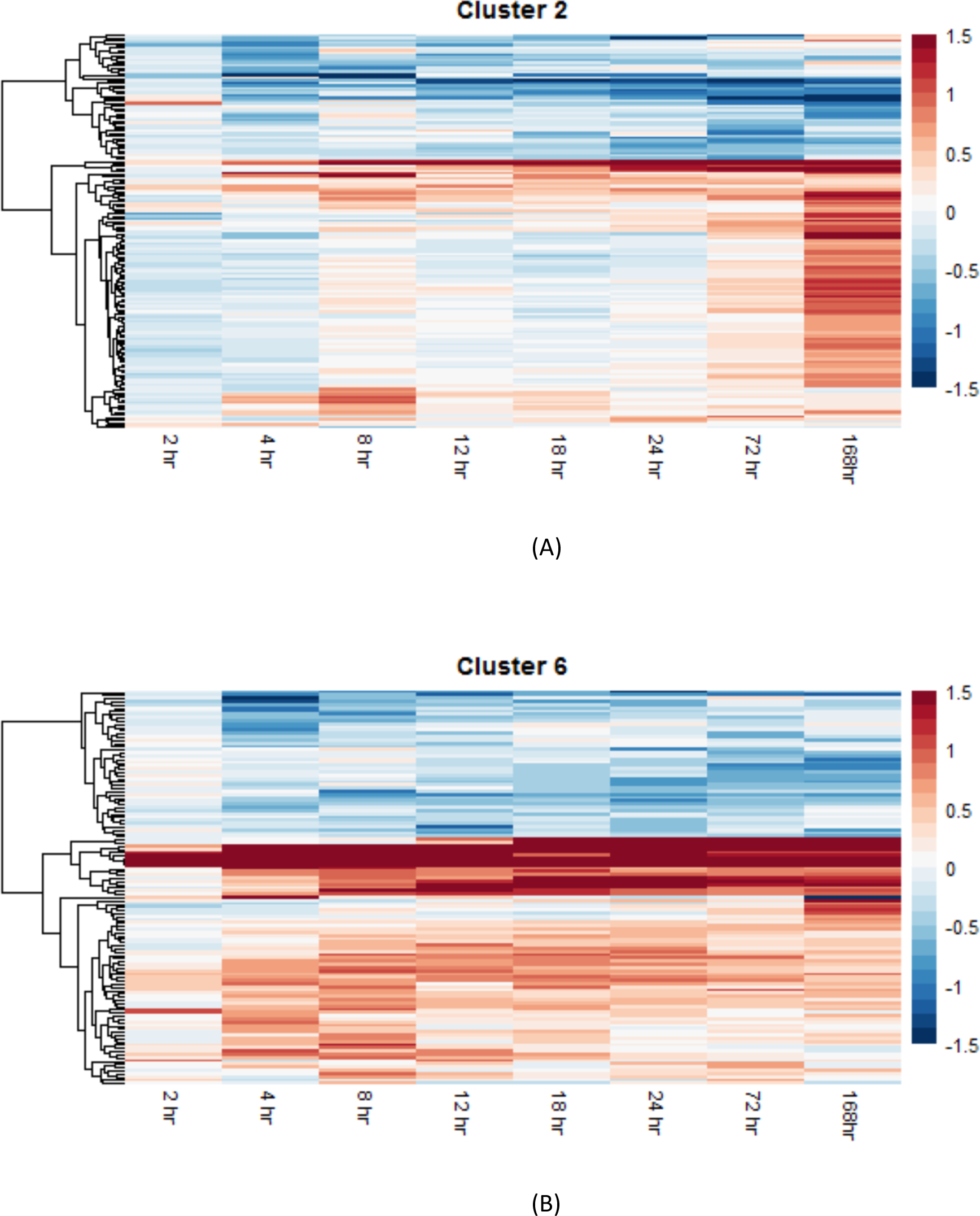
Heatmaps showing time courses of logz fold change for all 157 genes in Cluster 2 (A) and all 123 genes in Cluster 6 (B). For visualization of the heatmap, log2 fold change values > 1.5 were set to 1.5 and values < -1.5 to -1.5. Blue indicates downregulation and red upregulation.

**Figure 6:**
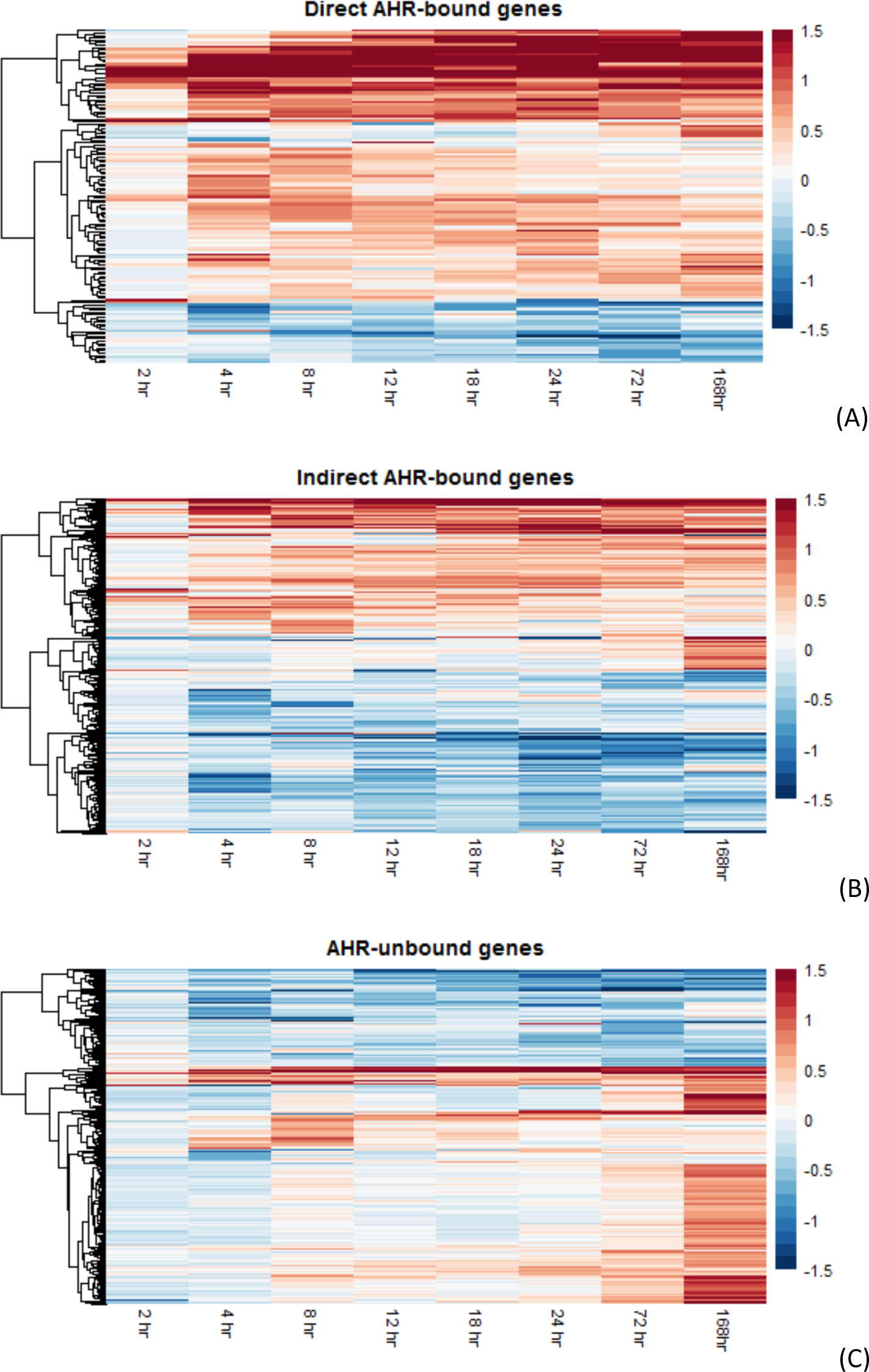
Heatmaps showing time courses of log2 fold change for 140 genes directly bound (A), 477 genes indirectly bound (B), and 574 genes unbound (C) by AHR. For visualization of the heatmap, log2 fold change values > 1.5 were set to 1.5 and values < -1.5 to -1.5. There are proportionately more upregulated genes in (A) compared to (B), and in (B) compared to (C). Blue indicates downregulation and red upregulation.

These differences between direct, indirect and unbound AHR target genes are also highlighted in overlaid box and violin plots (**Figure. 7A-D**), showing the respective distributions of expression level of the three groups of genes at multiple time points. At each time point shown, the middle 50% (first to third quartile) of the directly bound genes are all upregulated, while the indirectly bound group is symmetrically distributed with about half of the genes upregulated. In the unbound group, most genes are downregulated at earlier time points, but at 168 hrs., the distribution is considerably right-skewed with many genes upregulated. Overall, the directly regulated group has the highest median expression (except at 168 hrs.), and also has the most outliers on the high expression end, the furthest outlier being the Cyp1a1 gene. Given that the Cyp1a1 gene has a high number of DREs in its proximal promoter region (Li et al. 2014), we examined the relationship among expression level and number of proximal promoter DREs for the direct target genes at various time points (**Figure. 8A-D**). There is an increase in mean expression level with increasing number of DREs, a trend that gets stronger at later time points. This is likely due to a larger number of AHR molecules binding to DREs in the promoter regions, leading to a higher degree of activation of proximal genes. This hypothesis is supported by previous findings that in the human liver, genes with more transcriptional regulators bound in their promoter regions were more highly expressed (Odom et al. 2006).

**Figure 7:**
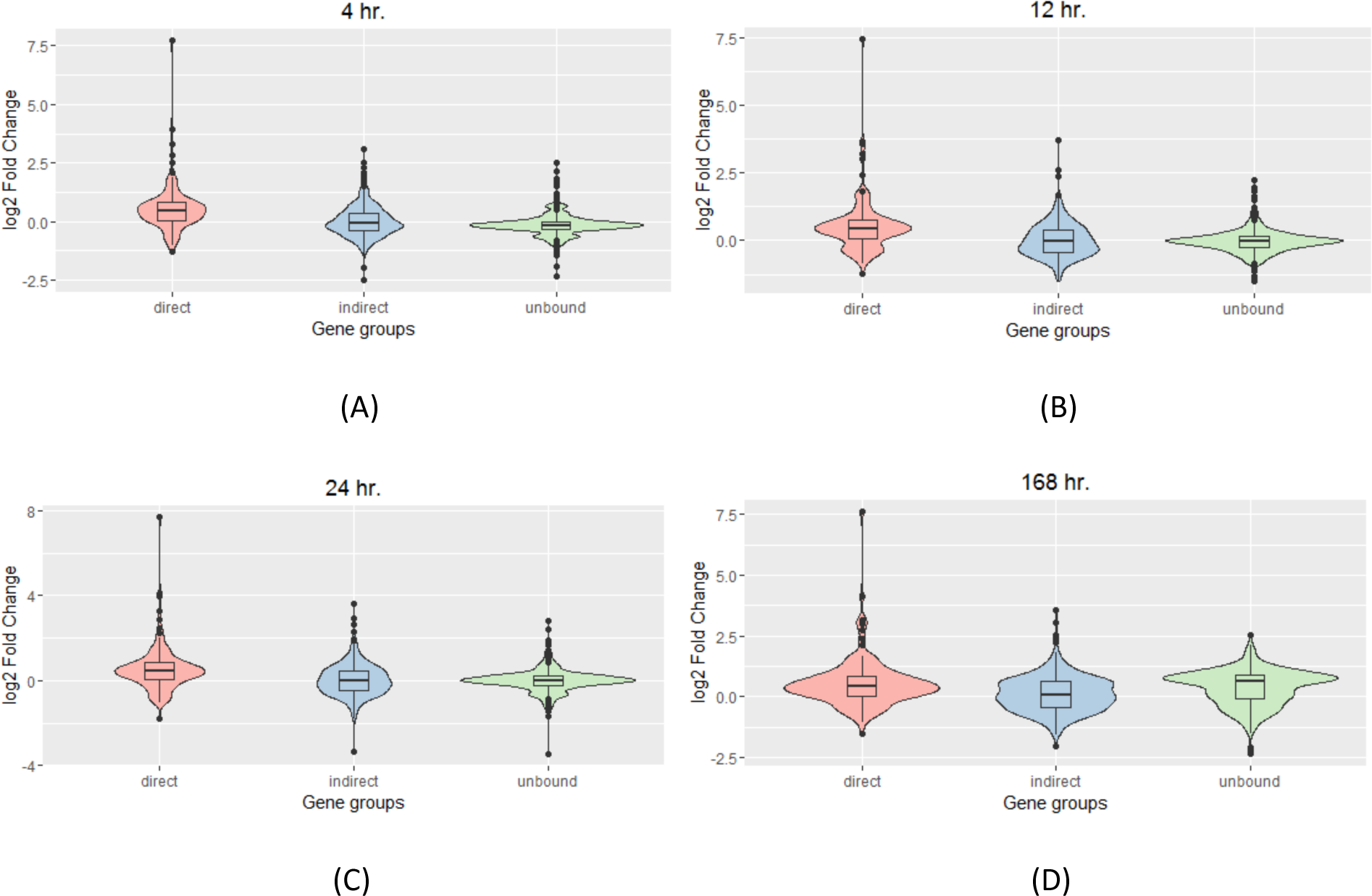
Overlaid box and violin plots showing the distribution in differential expression of direct (n = 140), indirect (n = 477) and unbound (n = 574) AHR target genes at 4 hr. (A), 12 hr. (B), 24 hr. (C) and 168 hr. (D). These plots illustrate the respective distributions of expression level of the three groups of genes at multiple time points, with the box plots illustrating the median, first and third quartile, and outliers; and the overlaid violin plots showing a rotated histogram of the distribution of gene expression.

**Figure 8:**
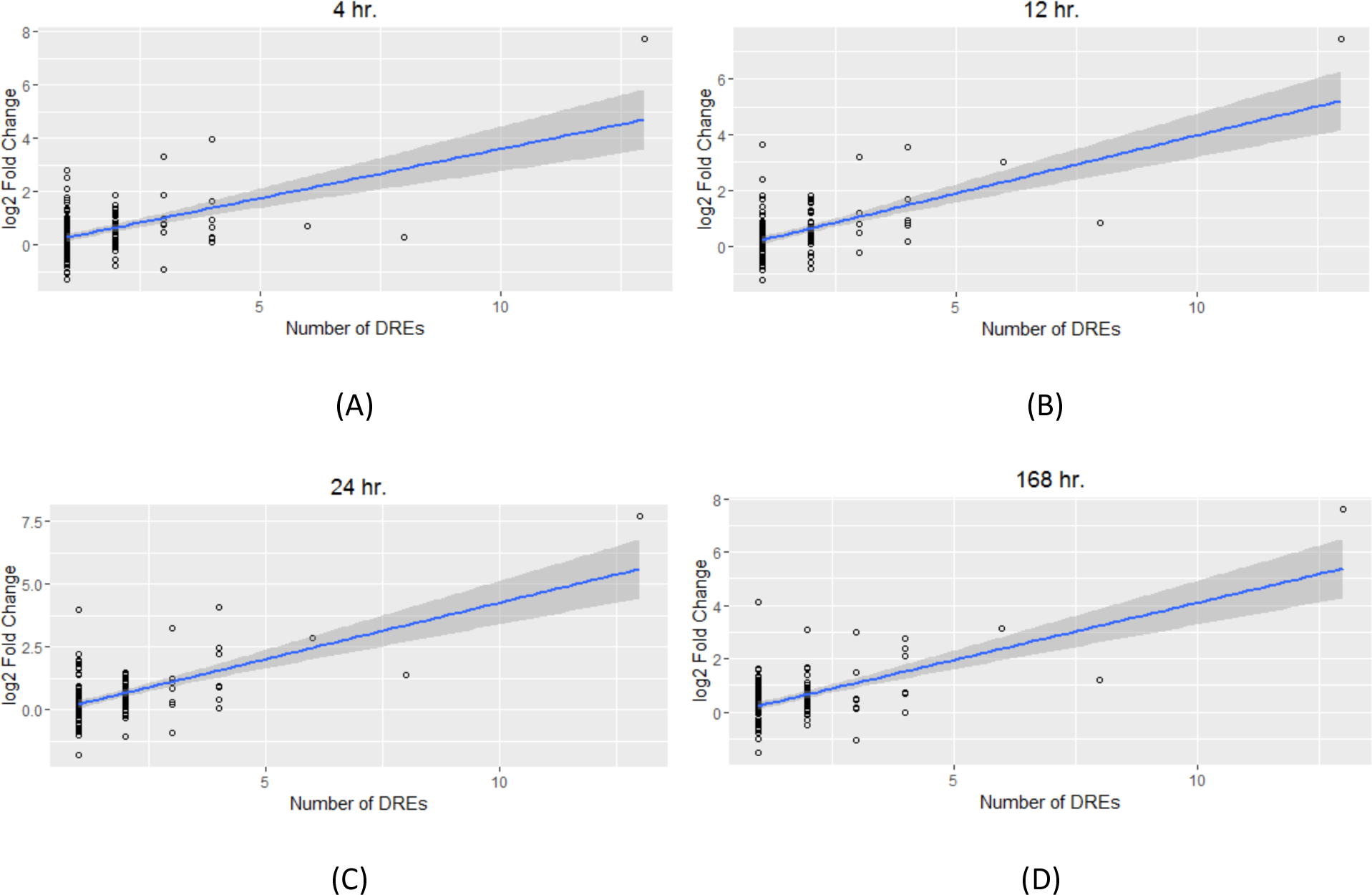
Increase in expression level of direct AHR target genes with number of DREs in proximal promoter regions at 4 hr. (A), 12 hr. (B), 24 hr. (C) and 168 hr. (D). Circles denote individual genes; linear regression fit shown in blue line with shaded region showing 95% confidence interval.

### Distinct gene clusters activate distinct biological processes

We carried out gene ontology (GO) analysis on the six major clusters of genes labeled in **Figure 4A**. The genes in the six clusters enrich for different groups of biological processes (**Figure 4B**). In particular, the genomic cluster (Cluster 6) is enriched for genes associated with metabolic processes and ribosome biogenesis, whereas the major GO categories associated with the non-genomic cluster (Cluster 2) are immune regulatory processes. Interestingly, Cluster 5 is enriched for cell migration and activation of cellular defense mechanisms. Presumably this reflects immune cell infiltration into the mouse liver under exposure to TCDD (Fader et al. 2015). Cluster 16 is also enriched for immune system response. Thus, co-regulated genes in the AHR network in the mouse liver show patterns of co-expression, and lead to differential downstream activation of biological processes.

## Discussion

Ligand-activated transcription factors underlie most major cellular response pathways. These TF-governed molecular pathways tend to have a similar organizational structure with key functional components that act as signal sensors (co-binding proteins) and transducers (protein kinases) to complement the central role of the TF (Simmons et al. 2009). The inactivated TF is typically sequestered in the cytoplasm or nucleus. Upon activation by its ligand (endogenous or exogenous molecule), the TF is able to bind specific response elements in the promoter regions of target genes and activate or inhibit expression of suites of genes in a coordinated manner. Beyond these “direct target” genes, there are additional genes that bind the master regulatory TF indirectly through tethering interactions with secondary TFs (George et al. 2011; Shen et al. 2011; McMullen et al. 2014). In fact, combinatorial control of gene expression by TFs is a common feature of cellular pathways, since binding sites are often clustered in the genome, allowing multiple TFs to act in a coordinated fashion to induce or suppress groups of genes in specific cell types under particular conditions (George et al. 2011). In addition, a surprisingly large number of genes are activated or inhibited in a “non-genomic” manner, showing no evidence of binding by the master regulatory TF of the stimulated pathway in their promoter regions (van der Meer et al. 2010; Dere et al. 2011b; Shen et al. 2011; McMullen et al. 2014). These observations collectively suggest that combining gene expression data from transcriptome profiling with high-throughput genome-wide analysis of TF binding can provide an integrated, systems-level view of the structure and function of transcription factor-governed molecular pathways (Blais and Dynlacht 2005; Walhout 2006; Dere et al. 2011b; Limonciel et al. 2015).

Accordingly, we have integrated TCDD-induced gene expression and multiple genome-wide TF binding data sets for a global view of the AHR regulatory pathway in the mouse liver. Using a combination of self-organizing maps and subspace clustering, we show that there is a pattern of co-regulated genes in the AHR pathway being co-expressed, as previously observed in *Saccharomyces cerevisiae* (Yu et al. 2003; Allocco et al. 2004). In particular, directly-bound, indirectly-bound and unbound AHR target genes have distinct patterns of gene expression, with the directly-bound group showing higher median expression. Further, among the direct AHR target genes, the expression level increases with the number of AHR-binding DRE sites in the proximal promoter regions. Finally, we found that co-regulated gene clusters activated distinct groups of downstream biological processes, with the AHR-bound genomic cluster enriched for metabolic processes and the AHR-unbound non-genomic cluster primarily activating immune processes. This work, together with other recent studies of the PPARα and estrogen receptor pathways (McMullen et al. 2014; Pendse et al. 2016), illustrates the application of bioinformatic and statistical tools for reconstruction and analysis of the transcriptional regulatory cascades underlying cellular stress response.

## Acknowledgments

The authors would like to thank Agnes (Forgacs) Karmaus, Arindam Banerjee, Rory Conolly and Qiang Zhang for helpful discussions. This work was supported by the US EPA STAR Program (EPA Grant Number: R835000) and the Superfund Research Program of the National Institute of Environmental Health Sciences (P42ES04911).

